# A normalized template matching method for improving spike detection in extracellular voltage recordings

**DOI:** 10.1101/445585

**Authors:** Keven J. Laboy-Juárez, Sei Ahn, Daniel E. Feldman

## Abstract

Spike sorting is the process of detecting and clustering action potential waveforms from extracellular voltage recordings to identify spikes of putative single neurons. Typically, spike detection is done using a fixed voltage threshold and shadow period, but this approach can lead to missed spikes during high firing rate epochs or noisy conditions. We developed a novel spike detection method utilizing a computationally simple form of template matching that efficiently detects spikes from candidate single units and is tolerant of high firing rates and electrical noise without a whitening filter. Template matching was based on a sliding cosine similarity between mean spike waveforms of candidate single units and the extracellular voltage signal. Performance was tested in whisker somatosensory cortex (S1) of anesthetized mice *in vivo*. The method consistently detected whisker-evoked spikes that were missed by a standard fixed voltage threshold. Detection was most improved for spikes evoked by strong stimuli (40-70% increase), less improved for weaker stimuli, and unchanged for spontaneous spiking. This reflected the failure of standard detection during spatiotemporally dense spiking. Template-based detection revealed higher signal-to-noise ratio for sensory responses and sharper sensory tuning. Thus, this template matching method (and other model-based spike detection methods) critically improve the quantification of single-unit spiking activity.

## Introduction

Extracellular single unit recording is widely used to monitor activity of neuronal populations, determine sensory tuning and evaluate stability and plasticity of neural coding over time^1, 2^. Because raw extracellular voltage signals incorporate spiking of many nearby neurons over background noise, spike sorting is used to isolate action potentials and assign them to putative single neurons via clustering^3-5^. Thus, high quality in both spike detection and clustering algorithms are critical to accurately measure single unit activity. Recent breakthroughs have significantly improved spike clustering procedures, greatly reducing the need of manual intervention and allowing recording of single unit activity in large neuronal populations over long periods of time^6-8^. Despite these advances, spike detection is still mostly done with a simple voltage threshold trigger, a method that can fail to detect spikes relative to background recording noise and during spatiotemporally dense population activity. Here we report an improved method for spike detection that aids in the accurate recovery of single unit spike trains, evaluated in the primary whisker somatosensory cortex of mice.

The standard spike detection procedure consists of setting a voltage threshold and classifying any voltage fluctuation that crosses this threshold as a putative spike. A brief segment of recorded voltage (termed a voltage clip; typically from ∼0.1 ms before to ∼1.4 ms after threshold crossing) is stored for later spike clustering. To prevent a single spike from being detected multiple times as separate events, a brief shadow period is enforced in which spike detection is disabled for ∼0.6 ms after the initial threshold crossing. Thus, the ability to detect spikes is directly determined by the voltage threshold and the shadow period. If the threshold is too stringent, low amplitude (yet potentially sortable) spikes will go undetected, which can cause entire single units to be missed, or cause the loss of a fraction of spikes from well-isolated single units whose mean spike amplitude is just above detection threshold.

Conversely, if detection threshold is too low, more noise and non-sortable spikes will cross threshold, increasing the fraction of time spent in shadow period, and thus reducing detection of larger, sortable spikes^9^. Despite the strong dependence of overall spike sorting on the voltage threshold, the value of the voltage threshold is often chosen arbitrarily, e.g. as a fixed multiple of the standard deviation of the voltage signal on a given recording channel.

The number of legitimate spikes that occur during the shadow period, and will therefore be missed in spike detection, depends on the temporal dynamics of local neuronal activity. Strongly correlated firing of nearby neurons on a short time scale and high temporal precision of neural responses will increase the number of shadowed spikes, as will greater overall mean firing rate. These conditions often occur in topographically organized brain regions, such as primary sensory cortex, when an optimal sensory stimulus for the local population is presented^10-12^ and are important for precise sensory feature representations^13-15^. As a result, more spikes are likely to be missed in response to locally preferred stimuli, which will distort measurements of neural tuning. To maximize spike detection under these conditions, an improved method of spike detection is needed. We developed a normalized template matching method (NTM spike detection) that uses a different type of threshold—waveform similarity to a candidate single-unit spike waveform, rather than spike amplitude, and applies a data-driven approach to optimize detection threshold for individual single units. We show that this approach reduces shadowed spikes and selectively increases detection of candidate spike waveforms relative to non-sortable spikes, yielding improved single-unit sensory responsiveness and tuning.

## Results

### The normalized-template-matching (NTM) method

Spike-sorting with the NTM method is a two-part process (**Fig. 1a**). First, an initial round of spike detection is performed using a standard fixed voltage threshold, and spike sorting is performed to cluster spike waveforms by their shape. The goal of this initial round is to identify mean spike waveforms of candidate, isolatable single-units. We term this initial round the ‘standard method’. This is followed by a second round where spike detection performed via the NTM template matching method, and the goal is to maximize spike detection for these specific candidate single units. Detected spikes are then clustered by spike sorting, as in the standard method. We use one particular spike clustering method here, but NTM can be coupled with any clustering method.

**Figure 1.**
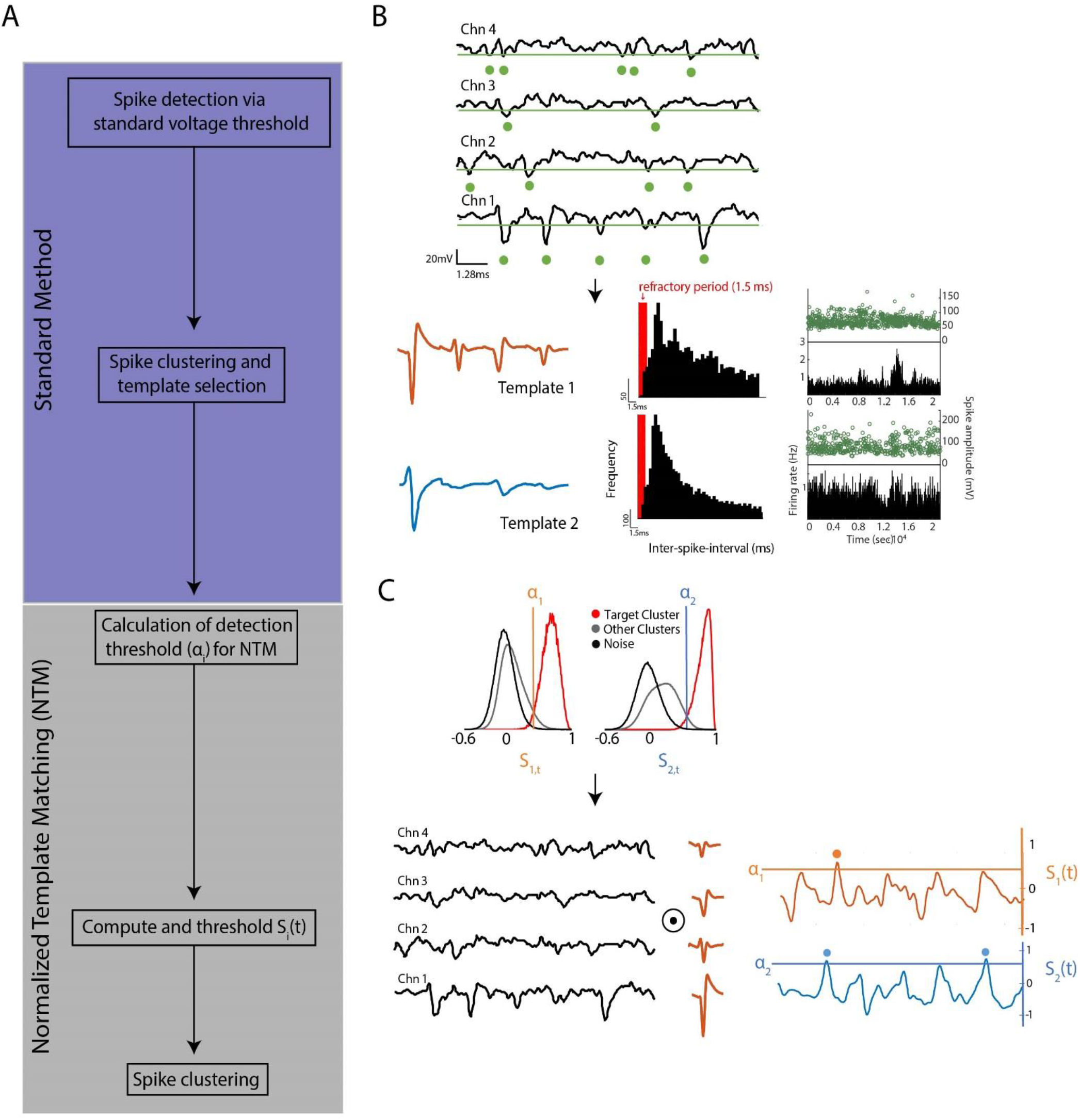
Normalized template matching algorithm for spike detection **(a)** Step-by-step description of normalized-template-matching (NTM). The standard method is an initial round of spike sorting with the usual voltage threshold trigger. NTM is the second round of spike sorting where NTM is used for spike detection. **(b)** Graphical schematic of the standard method. Top, segment of extracellular voltage signal. Green line is the fixed threshold (usually 3 standard deviations below the mean) and dots denote threshold crossing events. Bottom, mean spike waveform of 2 example isolatable single-units. ISI distributions and spike amplitude and firing rate as a function of time are also shown. **(c)** Graphical schematic of NTM. Top, calculation of detection threshold for each template based on the spikes detected in the standard method. S_i, t_ is the cosine similarity between the template and a spike. Target cluster are spikes that were assigned to the single-unit of interest. Bottom shows the calculation of the scaled cross-correlation S_i_(t) for each template. Dots denote events that were classified as spikes.

### First step: the standard method

In initial spike detection procedure, we identified spikes by applying a fixed threshold to band-pass filtered voltage traces that were referenced to a common average across recording channels^16^ (see Methods). Any negative-going voltage transient that crossed the chosen threshold (usually 3 standard deviations below the mean) and did not fall in a shadow period was classified as a spike (**Fig. 1b**). A shadow period of 0.66 ms was enforced after each threshold crossing. This method is only sensitive to voltage amplitude not shape; background recording noise and spikes from non-isolatable units (multiunit activity) can also pass threshold and be detected. Because a shadow period was used, both noise and multi-unit (non-sortable) activity will suppress detection of single-unit spikes.

After detecting spikes, a semi-automated clustering algorithm was used to sort detected spike waveforms into clusters (putative single-units). We used an open source toolkit that performed the threshold-based spike detection and clustered clipped spike waveforms with a hierarchical clustering algorithm^17, 18^. Briefly, clipped spikes were aligned with respect to their peak and over-clustered through recursive bisection. These clusters were then aggregated based on spike waveform similarity and inter-spike-intervals (ISI) while minimizing refractory period violations that are commonly assumed for single units. Final clusters were classified as isolatable single units by manual inspection using commonly used quality metrics^9^. These metrics were refractory period violations <0.5% of the total number of waveforms in the cluster, and < 30% of spikes missing because their amplitude did not exceed the user-defined voltage threshold (estimated from a Gaussian fit of the spike amplitude distribution; see Methods). These clusters were then used as candidate isolatable single units in the second step.

### Second step: NTM and re-clustering

We performed a second round of spike detection and sorting in which NTM was used to maximize spike detection for the candidate units identified in the first step. NTM detects spikes based on spike waveform similarity, not amplitude or absolute voltage threshold crossing. Waveform similarity is determined by calculating the cross-correlation between the extracellular voltage signal and the mean spike waveform (template) of each candidate single unit (Fig. 1c). We define the template for single unit *i* as μ_i_ = [μ_i,1_,μ_i,2_, …,μ_i, N_] where μ_i, c_ is the mean spike waveform of single unit *i* in electrode (or channel) *c*. The waveform μ_i, c_ is a vector with *L* samples with *L* = f_s_k, where f_s_ is the sampling frequency and k is the user-defined time window of voltage over which the spike waveform is defined (in this study k = 0.0015 s; from 0.1 ms before to 1.4 ms after threshold crossing). Similarly, we define the extracellular voltage signal as V(t) = [**v(t)_1_, v(t)_2_**,…, **v(t)_N_**], where **v(t)_c_** is a segment of voltage signal on electrode *c* from sample *t* to sample *t* + *L*; i.e. **v(t)_c_** = [v(t)_c_, v(t+1)_c_,…, v(t+L-1)_c_]. The cross-correlation between the signal V(t) and template μ_i_ is given by the expression:

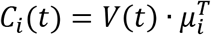

where (·)^T^ is the vector transpose. Thus the cross-correlation as a function of time is simply a sliding dot product between the voltage signal and the template. The dot product of these two vectors can be rewritten as:

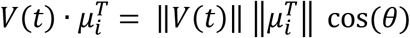

where ||·|| is the vector magnitude and θ is the angle between V(t) and μ_i_. Since ||μ_i_|| is constant in time, changes in C_i_(t) will only be due to changes in cos(θ) and ||V(t)||. Cosine of θ is a measure of similarity between the extracellular voltage segment and the template; a value close to 1 means that the two signals have a strong correlation in time (i.e. similar shape) because θ≈0. ||V(t)|| is directly related to the energy of the extracellular voltage segment and will increase during epochs of high-amplitude voltage fluctuations. These high-amplitude voltage events can be caused by spiking of nearby neurons or electrical noise. Since the events of interest consist only of those with a similar shape to μ_i_ we remove the influence of the signal energy by scaling the cross-correlation C_i_(t) to:

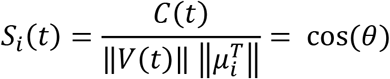

Thus, the scaled cross-correlation S_i_(t), which we refer to as normalized-template-matching (NTM), is equivalent to a sliding cosine similarity between V(t) and μ_i_ (Fig. 1c).

After computing S_i_(t) we calculate a threshold α_i_ such that any point in time where S_i_(t) ≥ α_i_ is detected as a spike and stored as a voltage clip for clustering. An important difference between NTM and the fixed voltage threshold method is that we compute ai through a data-driven approach rather than arbitrarily choosing a threshold relative to the noise floor. In this approach, we calculate the value of α_i_ through a receiver operating characteristic (ROC) curve such that the true positive detection rate (TP) of spikes from candidate units and correct rejections (CR) for multiunit activity and noise are maximized. For every single-unit *i* we compute the cosine similarity between each voltage clip and its template μ_i_:

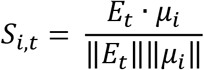

where E_t_ is a voltage segment that exceeded the amplitude threshold at time *t* and S_i, t_ is the cosine similarity between E_t_ and μ_i_. After spike clustering, the distributions of S_i, t_ for putative spikes that were classified as belonging to single unit *i* can be compared to the remaining putative spikes (**Fig. 1c**). The threshold α_i_ is chosen as the value of S_i, t_ that maximizes the percent correct classification of putative spikes belonging to single unit *i*, i.e. maximizes (TP + CR)/2. Importantly, voltage segments that don’t exceed the fixed voltage amplitude threshold (noise events) have a mean cosine similarity close to 0 that is separable from spikes from single units (**Fig. 1c**). This shows that the scaling operation prevents noise epochs from being detected as spikes. NTM thus represents a nonlinear filtering operation that is invariant to multiplicative scaling of the voltage signal and output is bounded to ±1. This framework is similar to two previous studies using linear filtering^19-21^ with the difference that NTM does not require the calculation and inversion of the noise covariance matrix (used for whitening the voltage signal).

### NTM improves spike detection

As previously mentioned, standard spike detection algorithms only use voltage amplitude to detect spikes and are thus agnostic to spike waveform shape. NTM restricts spike detection to spikes from candidate isolatable single-units (**Fig. 1b-c**). We tested whether this improves spike detection rates for single units in primary whisker somatosensory cortex (S1) of anesthetized mice during whisker stimulation. Recordings were made using multi-site silicon probes from layers 2/3 and 4, and spike detection and sorting were performed on groups of 4 nearby recording pads, treated as independent tetrodes. Recordings were common average referenced, filtered at .3-8 kHz and sampled at 31.25 kHz (see Methods).

**Fig. 2a** shows an example recording that illustrates how template matching can improve spike detection. Standard spike detection was performed using a fixed voltage threshold on each tetrode channel (red lines in A), followed by the 0.66-ms shadow period. Spike (iii) was successfully detected and found to be part of an isolatable cluster, whose mean waveform across the 4 tetrode channels is shown in the red-and-black inset graph. But spikes (i) and (ii) were censored by the shadow period of preceding voltage crossing events (triangles in inset), and thus not detected. Subsequent NTM for this candidate single unit 8 waveform detected all 3 spikes, and spike clustering determined that (i) and (ii) were part of the same single-unit cluster as spike (iii). Spike rasters and PSTHs for this unit revealed that NTM detected many more stimulus-evoked spikes following whisker deflections than the standard method (**Fig. 2b-c**). In contrast, spontaneous spiking (before stimulus onset) was largely similar between NTM and the standard spike detection method (**Fig. 2b-c**).

**Figure 2.**
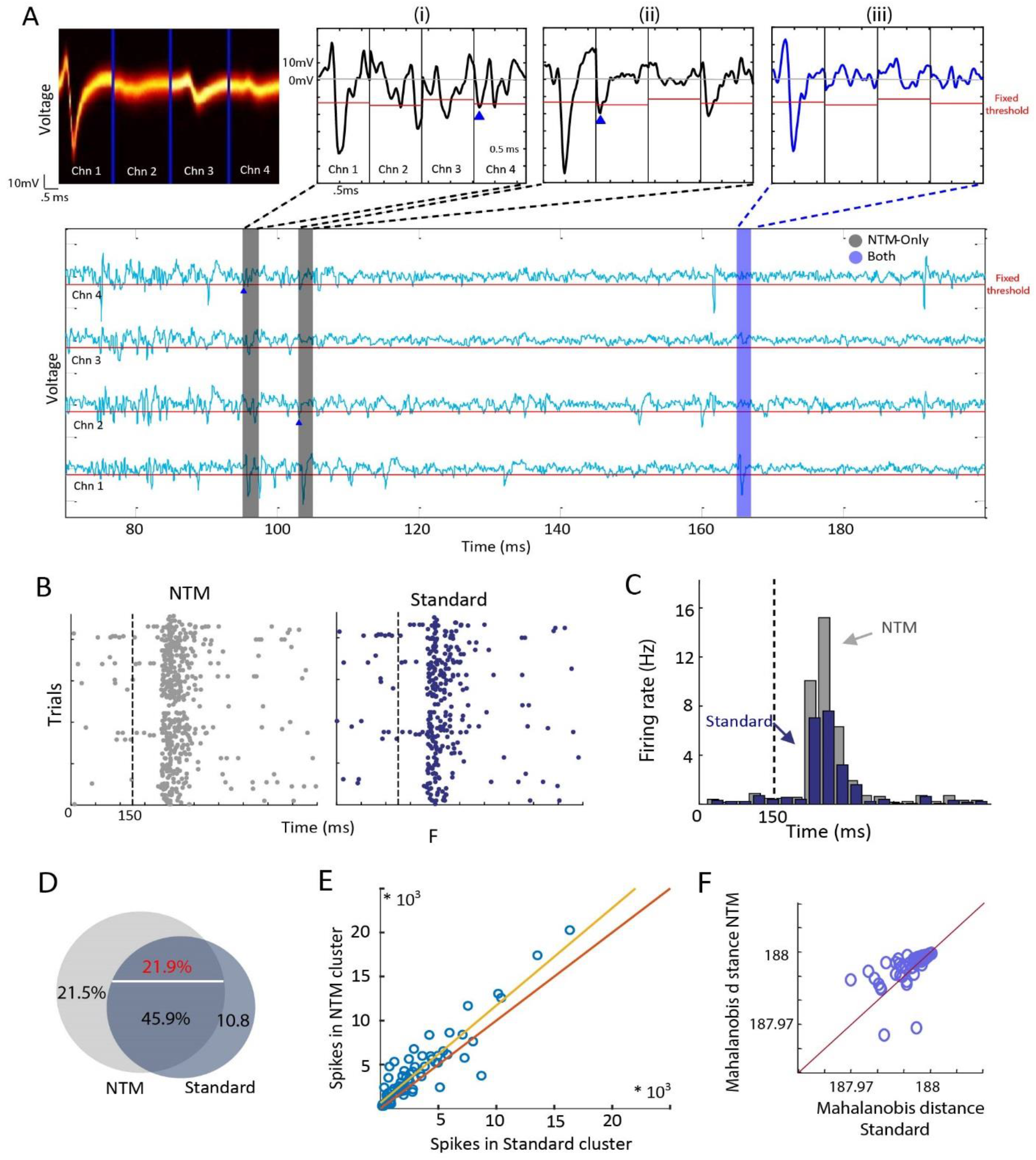
NTM detects more spikes than the standard voltage threshold trigger. **(a)** Example scenario where spikes from a single unit are missed by the voltage threshold trigger but detected by NTM. Top left, distribution of spike waveforms classified as belonging to an example single-unit. Bottom, segment of voltage signal from the 4 electrodes where the example single-unit was detected. Black and blue regions indicate spike events from the example single unit that were detected only with NTM and with both spike detection methods respectively. Top right show individual spike waveforms. Triangles are voltage threshold crossing events that suppressed spike detection. **(b)** Raster plot for an example single-unit using the standard and NTM spike detection methods. Black dash line is the onset of a whisker deflection. **(c)** PSTHs for the same example unit as (b) for NTM and standard spike detection. The standard PSTH was shifted in time for clarity. **(d)** Venn diagram showing the mean percentage of spikes only detected with NTM, standard voltage threshold or both. Percentage of spikes that were detected by both methods but assigned to different units (misclassified) after clustering is shown in red. **(e)** Total number of spikes detected for each single unit with NTM vs. standard voltage threshold. Misclassified spikes were not included. **(f)** Mean Mahalonobis distance for each single-unit with NTM and the standard voltage threshold.

We detected and sorted spikes from 28 tetrode recordings in 7 mice. 72 single units were found using a standard fixed voltage threshold, and then NTM was used to re-detect spikes from these candidate units. Across these single-units, the number of spikes detected after NTM was greater than those detected with the standard voltage threshold. On average, 21.48% of spikes detected by NTM were missed by the voltage threshold method. This is approximately twice the number of spikes that were missed by NTM but detected with the voltage threshold (**Fig. 2d**). In addition, 21.85% of spikes were detected by both methods but classified into different clusters after each step. This clustering error rate is within the expected range as assessed from simultaneous intracellular and extracellular recordings^22^ and is likely due to variability in automated clustering or manual curation procedures. Since this error is associated with clustering and not spike detection we discarded these spikes from subsequent analysis. NTM detected more spikes even after discarding misclassified spikes (**Fig. 2e**) (paired t-test p=2.18e-6, n=72 units; best-fit-line = 1.11x + 640.17). This suggested that NTM is better suited for measuring spiking activity.

Cluster quality after NTM was largely equivalent to the standard method, indicating that the additional spikes identified by NTM were similar in shape and amplitude to spikes identified by voltage threshold. To quantify this, we calculated the Mahalanobis distance of each spike to its cluster mean (spike waveform template), for spikes identified by standard voltage threshold and then subsequently by NTM.

For each cluster, we compared the mean Mahalanobis distance across all spikes before and after NTM. These were not significantly different (*p* = .3178) (**Fig. 2f**).

### Mean spike waveform shape is conserved after NTM

The greater number of spikes assigned to each cluster after NTM suggests that NTM can detect spikes that were missed by the voltage threshold. If these ‘new’ spikes are not due to clustering errors then the mean spike waveform of each single-unit should be relatively unchanged after NTM. **Fig. 3a** shows the mean waveform of two example single units before and after NTM, which have very similar overall waveforms. We quantified spike similarity by examining the spike amplitude at maximum negativity (reflecting the local current sink), positive peak (reflecting local repolarization) and spike width of the mean cluster waveform (**Fig. 3b**). We compared these three features for single-unit clusters before and after NTM. Across the population of 72 single units, no significant differences were found between NTM and standard method in negative spike amplitude, positive peak or spike width (paired t-test *p* =.38, .63 and .56 respectively), and the Pearson correlation was close to unity, with *r* = 0.93, 0.96 and .98 respectively (**Fig. 3c**). Thus, mean spike shape is not altered by NTM.

**Figure 3.**
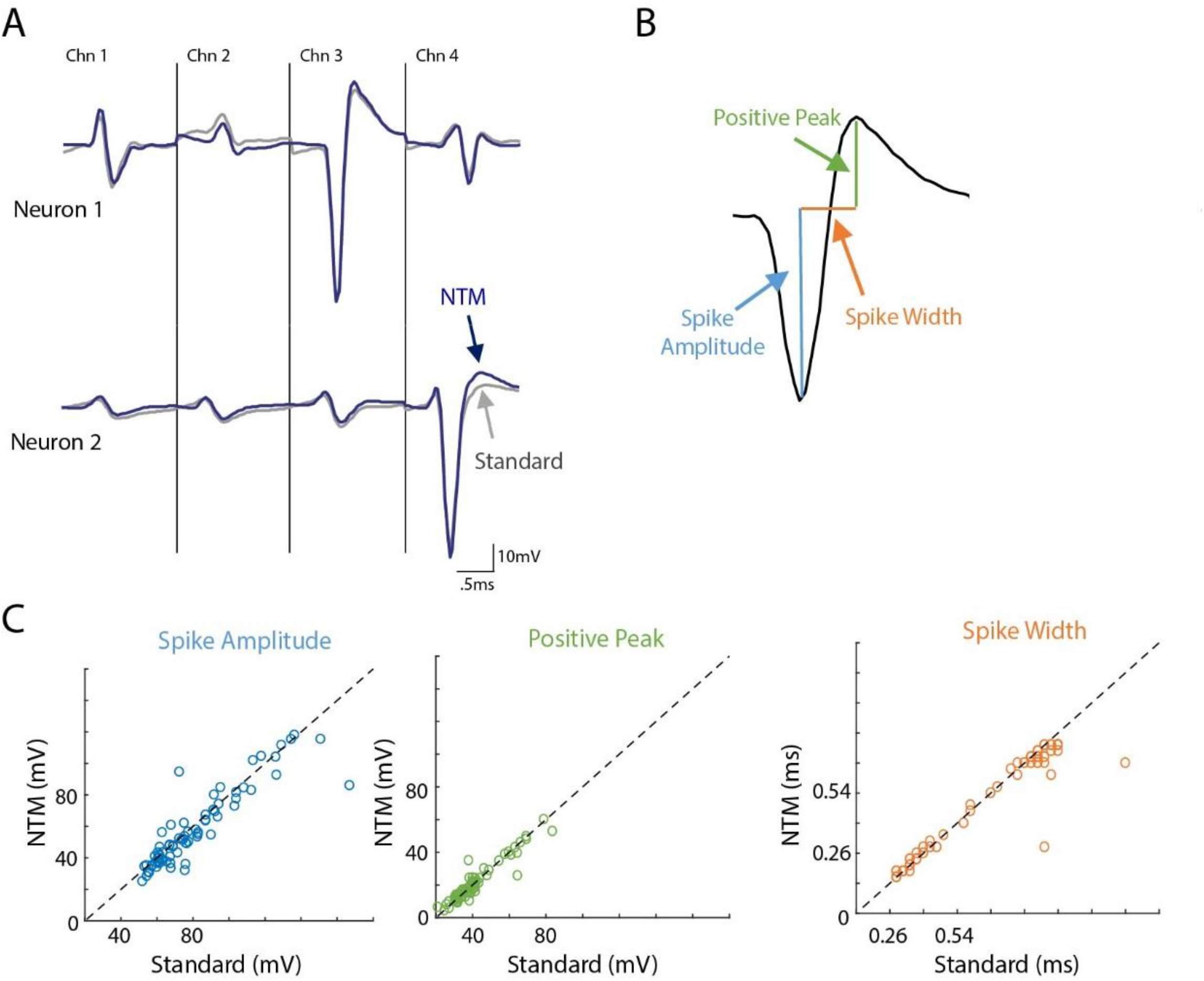
Mean spike waveform shape is conserved after NTM. **(a)** Mean spike waveforms of two example single-units with NTM and standard spike detection. **(b)** Schematic of the calculation of spike amplitude (current sink), positive peak (current source) and spike width (time difference between current sink and source). **(c)** Comparison across units for three spike waveform features.

### NTM improves measures of sensory responsiveness and sensory tuning

Dense local population activity can cause failure to detect legitimate spikes (false negatives or type II errors) because of shadow periods using the standard voltage threshold method. This raises a critical concern that spiking evoked by the strongest stimuli will be systematically under-estimated, thus reducing measures of sensory responsiveness and artifactually flattening sensory tuning curves. In S1, most neurons within a single cortical column spike most strongly to one facial whisker that is anatomically matched to that column (termed the columnar whisker, CW) ^23, 24^ This topographical organization means that CW deflections will elicit more spikes than deflections of surrounding whiskers (SWs), raising the possibility that type II errors are more common during CW deflections. This problem may be more acute in layer (L)4, which has higher neuronal density and stronger multi-unit population responses than in L2/3. We tested whether improvements in spike detection from NTM were systematically different across whisker stimuli and across layers.

For each single unit cluster (n=72), we compared the number of CW-evoked spikes (defined as spikes within 125 ms of stimulus onset) detected with standard and NTM methods. NTM detected significantly more CW-evoked spikes (paired t-test *p* = .0495) (**Fig. 4a**). In the same experiments, spontaneous firing rate was measured following sham whisker deflections, which were randomly interleaved. Spontaneous firing rate after sham stimuli was not significantly different between NTM and standard methods (*p* = .3915) (**Fig. 4a**, bottom). Thus, sensory responsiveness (signal-to-noise ratio) was underestimated with the standard voltage threshold. On average, CW-evoked firing rates were significantly greater with NTM for both layer 2/3 (2-way ANOVA, *p* = 1.9e-9) and layer 4 *(p* = 5.2e-7) populations (**Fig. 4b**). However, the greatest gain in detected spikes occurred for CW-over SW-evoked spikes, and for L4 over L2/3 (**Fig. 4c**). With NTM, layer 2/3 and layer 4 units had a 45.2% and 77.8% average gain in detected CW-evoked spikes respectively vs. a modest 13.4% and 11.1% average gain for SW-evoked spikes (this is average across 8 SWs). We calculated the whisker receptive field for each single unit after ranking SWs from strongest to weakest within each unit. The average whisker receptive field was sharper after NTM, because NTM improved spike detection most for the strongest stimuli (**Fig. 4d**). Thus, NTM dramatically enhanced detection of whisker-evoked spiking responses, especially for stimuli evoking particularly strong spiking responses across nearby neurons.

**Figure 4.**
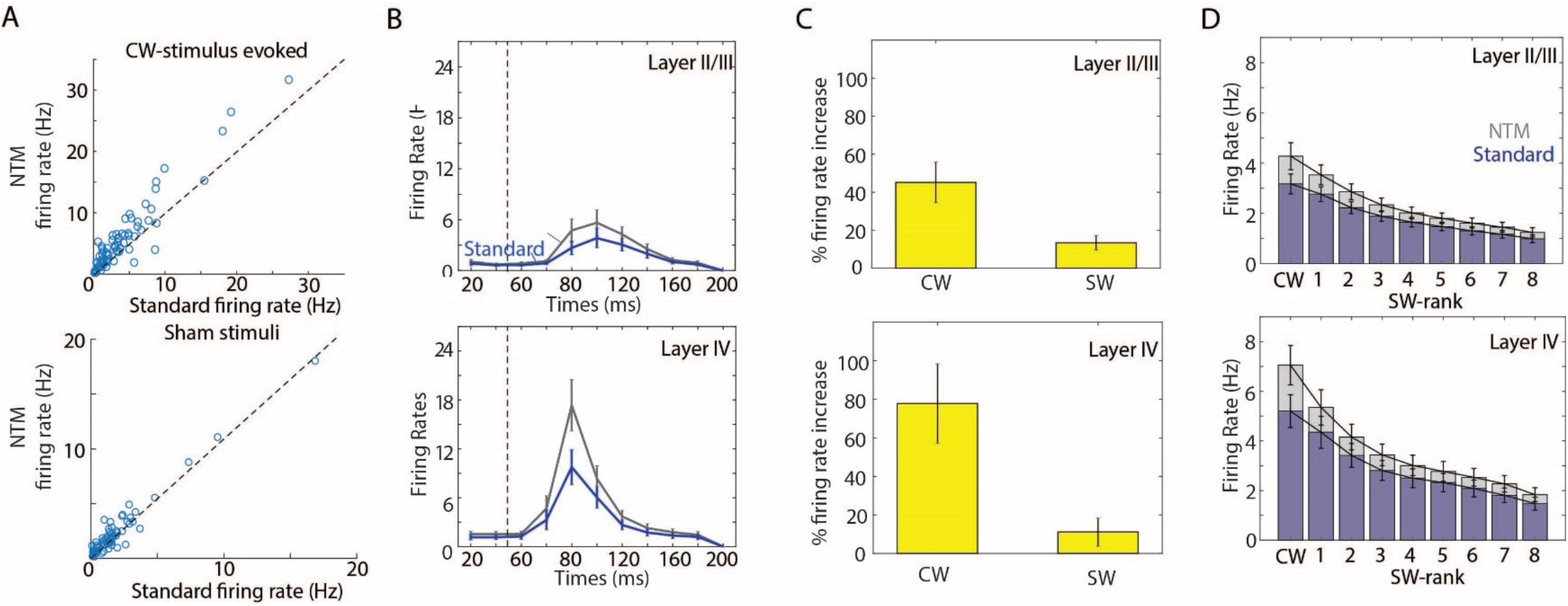
NTM improvements in spike detection are stimulus specific. **(a)** Top, comparison of measured CW-stimulus evoked firing rates with NTM vs. the voltage threshold trigger across all single-units. Bottom, same but for sham stimulation (i.e. spontaneous firing rates). **(b)** Average CW-stimulus PSTHs across layer II/III and layer IV single-units with NTM and the voltage threshold trigger. **(c)** Mean percent gain in measured CW- and SW-stimulus evoked firing rates with NTM relative to the standard voltage trigger. **(d)** CW and SW responses ranked by response strength for both NTM and the standard voltage threshold trigger.

### Source of newly detected NTM spikes

Spike events can be missed by the standard voltage threshold method for two reasons. First, the voltage amplitude of some spike events can be less than the user-defined threshold. Second, spikes can fall within the shadow period of a previous threshold crossing event. We tested which of these issues was primarily responsible for increased spike detection with NTM, under our experimental conditions.

For each NTM-detected spike that was missed by the standard threshold method, we determined whether the miss was due to a shadow period, or to spike amplitude that did not exceed detection threshold. For both L2/3 and L4 populations, most missed spikes were due to censoring by a shadow period. Shadowed spikes represented 60% of missed spikes in L2/3 and 70% of missed spikes in L4. Thus, most of the spikes missed by the standard voltage threshold method are due to local, temporally dense spiking near detection threshold.

## Discussion

We developed a spike detection algorithm based on template matching and tested its performance on extracellular recordings from silicon probe electrodes in S1 *in vivo*. Whisker receptive fields of S1 neurons are well characterized^23, 24^, with the majority of neurons having their strongest response to the anatomically matched columnar whisker (CW). Thus CW deflections evoke spatiotemporally dense spiking, a situation where spike detection is particularly challenging. We found that NTM substantially increased the number of stimulus-evoked spikes detected after CW deflection. The greatest gain in detected spikes was observed in L4, where spiking probability and spike timing precision are greatest ^10, 25^. Most spikes missed by the standard voltage threshold were due to the shadow period enforced after preceding voltage threshold crossing events. Thus, detection by a simple voltage threshold can yield many type II errors in spike detection (missed spikes), which compromise signal-to-noise and sensory tuning measurements. Because these mostly reflect shadow periods from prior low-amplitude spikes or near-threshold recording noise, they could in principle be avoided by using electrodes that better isolate a smaller number of nearby units (i.e., have a smaller effective recording radius). However, many modern recording devices, like silicon probes, are designed to maximize the number of recorded units, and thus include substantial non-sortable, low-amplitude spikes and recording ‘hash’ when local neural populations are strongly activated. NTM provides a method for improving spike detection with these devices, under conditions when strong and temporally dense network activity exists. We show that this enables more accurate measurement of stimulus-evoked spikes and sensory tuning.

NTM and its data-driven approach for choosing detection thresholds from template-filtered voltage traces is similar to some prior spike detection methods^19, 20^. The key improvement over these prior methods is that NTM does not require computing the noise covariance matrix and its inverse, which are computationally intensive steps. NTM computes a sliding cosine similarity rather than the cross-correlation between the whitened voltage signal and template, making NTM computationally and mathematically simpler. This also allows NTM to be easily implemented in applications that perform template matching in real-time (e.g. CED Spike2 and Ref^26^). We did not directly compare NTM performance with prior template based methods in this study.

Comparison of spike detection and sorting performance to ground truth requires simultaneous intra- and extracellular recordings of single-units, which we did not attempt here. Nevertheless, other groups have reported improved spike detection rates using other template matching methods relative to standard spike detection methods^19-21, 26^. Our data are consistent with this conclusion. Overall, our results strongly indicate that template-based spike detection methods such as NTM can greatly improve the quantification of single-unit spiking activity, especially in topographically organized areas like primary sensory cortex.

## Methods

### Surgical preparation and *in vivo* electrode placement

All procedures were approved by the UC Berkeley Animal Care and Use Committee and meet NIH guidelines. Male C57BL/6 mice (age: P28-45) used. Mice were anaesthetized with urethane and chlorproxithene (1.3 g/kg and 0.02 mg in 10 mL saline). Body temperature was maintained at 36.5° C using a feedback-controlled heating pad (FHC, 40-90-8D). Anesthetic depth was assessed via toe pinch and supplemental urethane (10% of initial dose) was provided as needed. The skull was exposed, cleaned and a stainless steel head-post was implanted. A 2 mm craniotomy was made over S1 (coordinates: 1.5 mm rostral, 3.3 mm lateral of bregma). The target column (the C1 or D1 whisker column) was localized using receptive field mapping of multi-unit activity in L4, recorded with a tungsten microelectrode.

A silicon laminar probe (NeuroNexus, 32 channel, 1 shank, poly2 or poly3 channel geometries (A1×32-Poly2-5mm-50s-177-A16 and A1×32-Poly3-6mm-50-177-A32) was then inserted radially into the target column via a small durotomy. The probe was slowly advanced until the deepest recording pad was in L4. Simultaneous L2/3 and L4 recordings were made at this depth. L2/3 and L4 were defined by microdrive depths as 100-417 and 418-587 μm below the pia^27^.

For all recording penetrations, we confirmed post-hoc that silicon probe multi-unit activity in L4 was tuned to C1 or D1, based on single-whisker deflections. In a subset of cases, recording location was confirmed by coating the recording electrode with DiI, perfusing the mouse, and recovering DiI staining in cytochrome oxidase stained flattened tangential sections, which show the L4 barrels.

### Whisker stimulation

Calibrated deflections were applied independently to a 3 × 3 grid of whiskers, centered on the columnar whisker for the recorded column. Stimuli were controlled using custom software in Igor Pro (Wavemetrics). Each whisker was trimmed to 8 mm length, and inserted into a glass tube carried on a piezoelectric bender actuator. The piezo was positioned to deflect the whisker at 5 mm distance from the face. Each whisker was deflected rostrocaudally with triphasic waveform that was shown previously to optimally drive S1 neurons and captured most of the evoked response variance in S1 (first common filter)^28^. The waveform was 40 ms duration, 300 μm peak amplitude, and had a mean frequency content of 53Hz.

The stimulus set consisted of 18 single-whisker deflections (9 whiskers with peak deflection amplitude in either the rostral or caudal direction) and additional 2-whisker combination stimuli that were not analyzed here. All stimuli were randomly interleaved at 0.6 s inter-stimulus interval, yielding an overall average deflection rate for any whisker of 3 Hz. Each recording was 3.5-6 hours. Sham stimuli (blank trials in which no whisker was deflected) were randomly interleaved with other whisker stimuli to quantify spontaneous spiking.

### Data acquisition and preprocessing

Recordings were amplified and bandpass filtered (Plexon Instruments PBX2/16sp-G50, × 1,000 amplification, 0.3-8 kHz bandpass) and digitized at 31.25 kHz. Noise was reduced by common average referencing^16^; the average voltage signal across all electrodes in the brain was calculated and subtracted from the signal in each electrode. Poly 2 electrode sites were divided into groups of 4 spatially adjacent channels (tetrodes; adjacent sites were 50μm apart). Poly 3 electrode sites were also divided into 4-channel tetrode groups, selecting channels the maximal signal-to-noise ratios located within a 50μm depth range. Spike detection and sorting were performed on each tetrode separately. For each electrode site, signal-to-noise was calculated as the kurtosis of the voltage signal.

### Spike sorting

In the standard method negative-going spikes were detected using an amplitude threshold (2.8-3.2 s.d. of noise floor), with a shadow period of 0.66 ms after each threshold-crossing. Detected spikes were then clipped (1.5-ms waveforms) for clustering and sorting. Isolatable single-units were labeled through manual inspection and had to satisfy the following criteria: <0.5% refractory period violations (defined as inter-spike interval < 1.5ms) and <30% estimated missed spikes (based on Gaussian fit of detected spike amplitudes relative to the voltage amplitude detection threshold). The mean spike waveform of each single-unit was then used as a template for NTM spike detection. As in the standard method, a shadow period was enforced after each voltage segment was detected as a spike (0.66 ms). Clustering was then redone as in the standard method.

All of the spike sorting used the open-access software UltraMegaSort2000^18^ (ums2k), implemented in Matlab. In the standard method spike detection, alignment, clustering and manual curation were all done with the functions provided in the ums2k package. For NTM, spike detection was done with custom software in Matlab (the NTM algorithm is described in detail in the results section) and spike alignment, clustering and manual curation were all done with the ums2k software package. Note that NTM can be implemented with any clustering algorithm.

## Code availability

All the code needed for performing the NTM method will be made available upon request via email to the corresponding author.

## Acknowledgements

This work was supported by NIH 1R37 NS092367 (to D.E.F.). KJLJ was supported by a NSF Graduate Research Fellowship and a NIH F99/K00 transition award.

## Author Contributions

KJLJ: Designed the normalized-template-matching algorithm and performed experiments.

SA: Analyzed data.

DEF: Project supervision.

KJLJ, SA and DEF: Wrote paper.

